# Addressing the digital divide in contemporary biology: Lessons from teaching UNIX

**DOI:** 10.1101/122424

**Authors:** Serghei Mangul, Lana S. Martin, Alexander Hoffmann, Matteo Pellegrini, Eleazar Eskin

## Abstract

Researchers in the biomedical sciences increasingly rely on applications that lack a graphical interface and require inputting code that, such as UNIX. Scientists who are not trained in computer science face an enormous challenge in analyzing the high-throughput data their research groups generate. We present a training model for use of command-line tools when the learner has little to no prior knowledge of UNIX.

## Main Text

The increasing amount of data generated by high-throughput genomics is reshaping the landscape of contemporary biomedical research into a data science (Markowetz 2017; Severin 2011; Spreafico et al. 2015). More biomedical research is performed *in silico* (Stevens 2013), and biomedical researches increasingly rely on applications that require inputting code or negotiating systems that lack a graphical interface (GUI), generally referred to as a UNIX command line interface (CLI). Applying computational techniques to analyze high-throughput data represents an enormous challenge to scientists not trained in computation, because these individuals must overcome the digital barrier and switch from GUI to CLI (Price 2012). As a result, major biomedical research institutions are challenged with supporting the analysis of genomic and other large-scale data generated by groups who traditionally have not received computational training (Miller and Alben 2012; Schneider et al. 2010).

Here we propose a model for addressing the digital divide in contemporary biology. Our approach helps biomedical researchers transition from using a GUI (e.g., M.S. Excel) to UNIX command line. The CLI is powerful, flexible, and allows greater control over bioinformatics analyses. In fact, the vast majority of bioinformatics software is now designed for the UNIX CLI (Altschul et al. 2013; Seemann 2013). Today’s big-data projects require that these researchers either learn how to use command-line tools or out-source their data analysis. Active engagement in analyzing the data they generate is essential to advancing interdisciplinary research in the biological sciences. However, biologists and medical researchers often lack formal training in the use of UNIX CLI. These research teams face several unique challenges and opportunities.

One approach is for biomedical researchers to delegate large-scale data analyses to bioinformatics cores. However, we believe that outsourcing analyses presents several problems for the biomedical researchers themselves. First, complex issues that arise during the analysis of genomic data are difficult to predict in advance. Projects often require much more effort than anticipated by research groups, leading core groups to struggle with insufficient funds to cover the time spent on analyses. Second, research groups utilizing the core often want to move the project in different directions from what was originally proposed. In the long term, exploring additional aspects of data can be inefficient for biomedical researchers when data analysis is delegated to a core group on an as-needed basis.

Another approach is to develop a GUI, such as Galaxy, that allows researchers with limited computational background to easily create, run, and troubleshoot analytical pipelines (Weber et al. 2017). While useful, providing researchers an alternative interface for command-line tools also has several drawbacks. These interfaces are more limited in computational power, and the graphical interface inherently limits the researcher’s flexibility in analyzing data.

To overcome these limitations, we believe that research groups should receive training and resources to analyze the data that they generate. This “training and collaboration” model encourages research groups to efficiently complete projects and advance their own skills. However, biological researchers without a background in computer science are often intimidated by applications that require inputting code or negotiating systems that lack a graphical interface, such as UNIX.

Traditional educational models for biological and medical researchers at the undergraduate and graduate level do not include computational training. Learning command-line tools as an advanced scholar is a challenge, because university computer science courses are designed for undergraduate students who enroll in an intensive, multi-year curriculum. Introductory-level computer science courses build the learners’ knowledge incrementally, are time-consuming, and are scheduled inflexible during the academic year. Therefore, there is growing demand for bioinformatics training in data or statistical analysis and interpretation skills, particularly in the format of dedicated small-group workshops led by a skilled trainer (Brazas et al. 2017).

Under this framework, we developed a three-day series of workshops that train students with no prior computational background to use UNIX CLI for analytical tasks. Our approach is geared for medical and biological researchers with no prior computational background and helps participants learn key commands and develop fundamental skills. The goal of these workshops is for first-time learners to acquire just enough knowledge and skills to independently use small, yet powerful, command-line commands for rapid exploration and modification of data.

Over a span of six years, post-doctoral scholars affiliated with the Collaboratory of the Institute for Quantitative and Computational Biosciences (QCBio) at the University of California, Los Angeles (UCLA) have taught these workshops to over 400 people. We engage undergraduates, graduate students, postdoctoral scholars, and faculty from biological and medical disciplines in groups of approximately 15 to 20 individuals.

Qualitative feedback has been overwhelmingly positive; after nine hours of instruction, participants report that they are able to effectively use UNIX CLI to manage and analyze their data. Many report mastering fundamental skills, such as directly entering functional commands line-by-line into a workbench that manages multiple platforms and a unified file system—without the familiar aid of a GUI.

We offer four specific suggestions for teaching the UNIX command line. First, we minimize jargon and discipline-specific technical terminology. When unavoidable, we introduce terminology with a clear definition and explanation of the term’s context and application. Second, we present concepts at a slow and incremental pace. First-time learners often advance in the workshop at different paces. We regularly pause the course to walk around the class and provide one-on-one tutoring. These individual sessions provide an opportunity for students to ask questions that they might otherwise not ask in front of the class. In our workshops, new concepts are introduced stepwise, building upon the previous concept. Thus, we view such interruptions as a way to guarantee that students acquire the necessary skills before moving on to the next unit.

Third, we introduce key terms metaphorically. Rather than cultivate a deep understanding of fundamental computer science principles, learners in our workshops can quickly assimilate introduced techniques and apply skills within the context of their research project. For example, the concept of a “variable” in computer science is highly technical and requires substantial knowledge of informatics in order to grasp. We introduce this term metaphorically; as in, “the variable is a box where you store the numbers.” Finally, we adopt a flexible pedagogy. We follow no set background philosophy; instead, we introduce fundamental concepts as needed and offer a substantial amount of hand-on examples and personal guidance to consolidate the learner’s newly-acquired knowledge.

We also developed an evaluation plan to continually improve the quality of our workshops. Before and after each workshop, we administer a quiz to assess the efficiency of the training model. Observational results suggest that our model can successfully train first-time users of command-line input systems to complete data analysis tasks. In one group, we observe complete elimination of the “little knowledge” category—all first-time learners moved from 0-70% to 25-100% after a short series of intensive workshops totaling nine instructional hours. We also see a shift of the entire cohort from including scores of 0% to scores of 25-100%.

We developed a publically available resource with workshop materials at: https://qcb.ucla.edu/collaboratory/workshops/collaboratory-workshop-1/

Training biomedical researchers in using UNIX to manage and analyze data appears to be successful with a series of workshops. The key challenge seems to lie not with inexperienced UNIX users’ aptitude for grasping analysis with CLI, but with the instructor’s ability to develop and teach curricula that unpacks computational skills in an approachable and digestible manner.

Our approach is easily reproduced and particularly useful for institutions where researchers from the biomedical sciences engage in big-data projects and frequently outsource computational analysis. An ability to analyze high-throughput data represents a competitive advantage for biological and medical researchers in today’s age of big data and next generation sequencing. UNIX is an “entry ticket” to bioinformatics; by gaining familiarity with UNIX, biomedical researchers may find it easier to engage with other applications and programming languages that are commonly used in computational biology.

We have also developed other workshops (n=15) that use similar teaching strategies. These workshops include, among others, “Intro to R and Bioconductor” and “Informatics for RNA-sequence Analysis.” Workshop materials of all workshops conducted through QCBio are publically available at:https://qcb.ucla.edu/collaboratory/workshops/.

